# Contribution of systemic and somatic factors to clinical response and resistance in urothelial cancer: an exploratory multi-omic analysis

**DOI:** 10.1101/086843

**Authors:** Alexandra Snyder, Tavi Nathanson, Samuel Funt, Arun Ahuja, Jacqueline Buros Novik, Matthew D. Hellmann, Eliza Chang, Bulent Arman Aksoy, Hikmat Al-Ahmadie, Erik Yusko, Marissa Vignali, Sharon Benzeno, Mariel Boyd, Meredith Moran, Gopa Iyer, Harlan S. Robins, Elaine R. Mardis, Taha Merghoub, Jeff Hammerbacher, Jonathan E. Rosenberg, Dean F. Bajorin

## Abstract

**Background:** Inhibition of programmed death-ligand one (PD-L1) with atezolizumab can induce durable clinical benefit (DCB) in patients with metastatic urothelial cancers, including complete remissions in patients with chemotherapy refractory disease. Although mutation load and PD-L1 immune cell (IC) staining have been associated with response, they lack sufficient sensitivity and specificity for clinical use. Thus, there is a need to evaluate the peripheral blood immune environment and to conduct detailed analyses of mutation load, predicted neoantigens and immune cellular infiltration in tumors to enhance our understanding of the biologic underpinnings of response and resistance.

**Methods and Findings:** The goals of this study were to (1) evaluate the association of mutation load and predicted neoantigen load with therapeutic benefit, and (2) determine whether intratumoral and peripheral blood T cell receptor (TCR) clonality inform clinical outcomes in urothelial carcinoma treated with atezolizumab. We hypothesized that an elevated mutation load in combination with T cell clonal dominance among intratumoral lymphocytes prior to treatment or among peripheral T cells after treatment would be associated with effective tumor control upon treatment with anti-PD-L1 therapy. We performed whole exome sequencing (WES), RNA sequencing (RNA-seq), and T cell receptor sequencing (TCR-seq) of pre-treatment tumor samples as well as TCR sequencing of matched, serially collected peripheral blood collected before and after treatment with atezolizumab. These parameters were assessed for correlation with DCB (defined as progression free survival (PFS) > 6 months), PFS, and overall survival (OS), both alone and in the context of clinical and intratumoral parameters known to be predictive of survival in this disease state.

Patients with DCB displayed a higher proportion of tumor infiltrating T lymphocytes (TIL) (n=24, Mann-Whitney p=0.047). Pre-treatment peripheral blood TCR clonality below the median was associated with improved PFS (n=29, log-rank p=0.048) and OS (n=29, log-rank p=0.011). Patients with DCB also demonstrated more substantial expansion of tumor-associated TCR clones in the peripheral blood 3 weeks after starting treatment (n=22, Mann-Whitney p=0.022). The combination of high pre-treatment peripheral blood TCR clonality with elevated PD-L1 IC staining in tumor tissue was strongly associated with poor clinical outcomes (n=10, HR (mean)=89.88, HR (median)=23.41, 95% CI (2.43, 506.94), p(HR>1)=0.0014). Marked variations in mutation loads were seen with different somatic variant calling methodologies, which in turn impacted associations with clinical outcomes. Missense mutation load, predicted neoantigen load and expressed neoantigen load did not demonstrate significant association with DCB (n=25, Mann-Whitney p=0.22, n=25, Mann-Whitney p=0.55, and n=25, Mann-Whitney p=0.29 respectively). Instead, we found evidence of time-varying effects of somatic mutation load on progression-free survival in this cohort (n=25, p=0.044). A limitation of our study is its small sample size (n=29), a subset of the patients treated on IMvigor 210 (NCT02108652). Given the number of exploratory analyses performed, we intend for these results to be hypothesis-generating.

**Conclusions:** These results demonstrate the complex nature of immune response to checkpoint blockade and the compelling need for greater interrogation and data integration of both host and tumor factors. Incorporating these variables in prospective studies will facilitate identification and treatment of resistant patients.

## Introduction

Atezolizumab has demonstrated responses in 15-25% of patients with advanced urothelial carcinoma and improved survival compared to historical expectations [1,2]. Similar to predictive factor analyses in melanoma, colon cancer and non-small cell lung cancer studies with other checkpoint blockade agents, Rosenberg and colleagues reported a statistically significant association between mutation load and response to atezolizumab in urothelial cancer patients [2]. However, mutation load in the atezolizumab study was predicted based on an estimate using a targeted panel and not with WES. Similar to findings from prior studies, the association between this predicted mutation load and outcomes in patients with urothelial cancer was not dichotomous; there were tumors from patients with elevated mutation load that did not respond to therapy, and vice versa. Additionally, positive PD-L1 staining of infiltrating immune cells by immunohistochemistry was associated with, but poorly predicted, response. A statistical model suggested that both PD-L1 staining and mutation load impacted the likelihood of response. However, the authors did not recommend their clinical use.

Collectively, studies to date imply that a combination of immune parameters are necessary to gain further precision in determining the likelihood of benefit from these immunotherapies and that a single biologic marker will be insufficient. There have been few attempts to integrate molecular and immunologic data from patients treated with checkpoint blockade and their tumors. Consequently we performed whole exome (WES), RNA, and T cell receptor (TCR) sequencing of tumor samples from patients treated with atezolizumab as well as TCR sequencing of matched, serially collected peripheral blood.

## Methods

### Ethics Statement

All research involving human participants was approved by the authors' Institutional Review Board (MSKCC IRB), and all clinical investigation was conducted according to the principles expressed in the Declaration of Helsinki. Written informed consent was obtained from the participants.

### Analysis plan

The analyses that were and were not included in the pre-specified analysis plan are detailed in S1 Pre-Specified Analysis Plan.

### Patients and clinical characteristics

All patients had locally advanced or metastatic urothelial carcinoma and were treated at Memorial Sloan Kettering Cancer Center (n=29) on protocol NCT02108652 [2]. All patients initiated therapy in 2014 and were treated with atezolizumab 1200 mg IV every 3 weeks, and provided written consent according to Institutional Review Board-approved protocols permitting tissue and blood collection, sequencing, and correlative studies. Patient tumor samples were assessed prospectively and centrally (by HistoGeneX, Brussels, Belgium) for PD-L1 expression by immunohistochemistry with the SP142 assay (Ventana, AZ, USA) [1]. The PD-L1 tumor-infiltrating immune cell (IC) status was defined by the percentage of PD-L1-positive immune cells in the tumor microenvironment: IC0 (<1%), IC1 (≥1% but <5%), and IC2/3 (≥5%) as defined in the original study. Six patients had multiple samples evaluated for PD-L1 IC status; for four of these patients, the sample used for PD-L1 IC status in this analysis was the same as the sample that was whole exome sequenced. For the remaining two, the PD-L1 status of one patient’s tumor (1994) used a separate primary tumor sample that also agreed in status with a metastatic sample; the other patient’s tumor (6229) used a metastatic sample site that agreed in status with another metastatic sample site. Smoking status was evaluated using previously completed self-reported smoking questionnaires or review of medical records. One patient was excluded because the patient did not consent to correlative studies beyond PD-L1 testing that was performed as part of the clinical trial.

### Tumor and blood samples

All tumor tissue used for sequencing was obtained prior to dosing with atezolizumab. Tumor samples used for whole exome sequencing were all formalin-fixed paraffin embedded (FFPE). The presence of tumor tissue in the sequenced samples was confirmed by examination of a representative hematoxylin and eosin-stained slide by a genitourinary pathologist (H.A.). Peripheral blood mononuclear cells (PBMCs) were isolated and stored as previously described [3]. PBMC were collected pre-treatment and during treatment.

### Clinical efficacy analysis

Tumor responses to atezolizumab were evaluated by CT scan every 9 weeks for the first 12 months following day 1 of cycle 1. After 12 months, tumor assessments were performed every 12 weeks. The response evaluation criteria in solid tumors (RECIST) version 1.1 was used to define objective clinical responses by the institutional radiologist.

### DNA extraction and high-throughput TCRβ sequencing

Tumor samples from patients 0522 and 6800 were excluded from tumor TCR analyses after failing quality control. Patients 0979, 7592, and 8214 did not have available tumor for TCRβ sequencing and were therefore excluded from tumor TCR analyses as well. Genomic DNA was purified from total PBMCs and tumor samples using the Qiagen DNeasy Blood extraction kit. The TCRβ CDR3 regions were amplified and sequenced using immunoSEQ® (Adaptive Biotechnologies, Seattle, WA) as previously described [4]. In brief, bias-controlled V and J gene primers were used to amplify rearranged V(D)J segments for high-throughput sequencing at ∼20X coverage. After correcting sequencing errors via a clustering algorithm, CDR3 segments were annotated according to the International ImMunoGeneTics Collaboration [5] to identify the V, D, and J genes that contributed to each rearrangement. A mixture of synthetic TCR analogs in each PCR was used to estimate the absolute template abundance (i.e., the number of cells bearing each unique TCR sequence) from sequencing data, as previously described [6]. The estimated TIL content was calculated as previously described [6−8]. To determine TIL content in FFPE samples as a T cell fraction, we amplified several housekeeping genes and quantitated their template counts to determine the amount of DNA usable for TCRβ sequencing. ImmunoSEQ then amplifies and sequences the molecules with rearranged TCRβ chains. Because the immunoSEQ assay aligns sequences to the IMGT database, sequences are annotated as complete VDJ rearrangements or non-productive rearrangements (a stop codon or out of frame CDR3 region was generated during VDJ recombination in one of the alleles); all downstream analysis in this work proceeded with complete, productive sequences. To estimate the number of starting templates that were in the sample, the number of sequence reads for each TCRβ sequences is measured. Synthetic control templates were also spiked into each sample, thereby enabling quantitation of input TCRβ templates from the read counts. For each sample, Shannon entropy was also calculated on the clonal abundance of all productive TCR sequences in the data set. Shannon entropy was normalized to the range by dividing Shannon entropy by the logarithm of the number of unique productive TCR sequences in the data set. This normalized entropy value was then inverted to produce the clonality metric. Those T cell clones whose frequencies differed between samples from a given subject taken at different time points, or between cell populations (e.g., between total PBMCs and tumor) were computationally identified as previously described [9]. The input data consisted of the absolute abundance for each TCR clone in each sample. Fisher’s exact test was used to compute a p-value for each clone across the two samples, against the null hypothesis that the population abundance of the clone is identical in the two samples. We corrected for multiple testing to control FDR using the Benjamini-Hochberg procedure and employed a significance threshold of 0.01 on adjusted p-values.

### Whole Exome Sequencing

Twenty-six FFPE-derived tumor and frozen PBMC-derived normal paired samples were sequenced by exome hybrid capture using the IDT xGen Whole Exome Panel (https://www.idtdna.com/pages/products/nextgen/target-capture/xgen-lockdown-panels/xgen-exome-panel) and standard protocols. Briefly, each sample was used to create a barcoded Illumina library, tumor samples were pooled at optimal multiplex to create an equimolar pool into the hybrid capture reaction, which was performed according to the manufacturer’s suggested protocol. Similarly, normal samples were pooled and introduced to the hybrid capture reaction. Following the recovery of captured library fragments, PCR amplification was performed, the resulting pools of fragments were quantitated using qPCR (Kapa Bio), and sequenced in separate lanes by paired end 150 bp reads, using the Illumina HiSeq 4000. Whole exome sequencing results for one sample (for patient 4072) were excluded after failing to meet coverage requirements.

### Somatic Variant Calling

DNA sequencing data for the tumor and normal samples were aligned to the GRCh37 reference using bwa-mem (v. 0.7.10) with default settings. The resulting BAMs were processed through Picard MarkDuplicates and the GATK (v. 3.5-0) pipeline including Base Quality Score Recalibration and Indel Realignment. Single nucleotide variants were called from Mutect (v. 1.1.6) and Strelka (v. 1.0.14) with default settings. Variants from either call were included and the variants calls were further filtered to those with depth (in normal and tumor samples) ≥ 7 reads, > 10% tumor VAF and ≤ 3% normal VAF [10]. Mutations per megabase was computed by normalizing the number of mutations by the number of exonic loci with ≥ 7 reads in normal and tumor samples, calculated using Pageant (https://github.com/hammerlab/pageant).

Variants were annotated as missense variants by Varcode (v. 0.5.10, https://github.com/hammerlab/varcode) and PyEnsembl (v. 1.0.3, https://github.com/hammerlab/pyensembl) using Ensembl Release 75 and annotated as deleterious using PolyPhen (v. 2.2.2). DNA damage response (DDR) genes were gathered from [11,12].

Mutational signatures were inferred from the somatic mutation calls using deconstructSigs (v 1.6.0).

### HLA typing

HLA types for each patient were computed from the normal sequencing data using OptiType (v. 1.0.0).

### RNA-seq

RNA was extracted from twenty-six FFPE tumor samples and evaluated for quality and quantity using the Agilent RNA pico chip. Each sample was prepared for sequencing by constructing an Illumina Tru-Seq Stranded RNA kit, according to the manufacturer’s protocol. The resulting libraries were amplified by PCR, quantitated, pooled and processed through a hybrid capture intermediate using the IDT xGen Exome reagent (as above). The captured fragments were quantitated, diluted and were sequenced using 2 x 150 bp paired end reads on the Illumina HiSeq 4000.

The RNA sequencing data were aligned to the GRCh37 reference an Ensembl Release 75 using STAR (v. 2.4.1d) and transcript quantification was performed using kallisto (v. 0.42.3). The STAR alignment was only used for identifying variant-supporting reads in the RNA. For gene-level analysis, the transcript quantifications were aggregated to the gene level using tximport (http://f1000research.com/articles/4-1521/v1).

Expressed mutations and neoantigens were computed using Isovar (v. 0.0.6, https://github.com/hammerlab/isovar), based on the RNA reads overlapping each mutation. Sleuth (v. 0.28.1) was used for differential expression analysis and GSEA was used for pathway enrichment analysis. ESTIMATE was used to quantify immune and stromal scores from RNA-seq data.

### Neoantigen calling

Neoantigens were computed from all nonsynonymous mutations using Topiary (v. 0.1.0, https://github.com/hammerlab/topiary) and NetMHCCons (v. 1.1) with HLA alleles calculated by OptiType. As with expressed mutations, expressed neoantigens were those supported in the RNA with at least 3 uniquely-mapped reads matching the cDNA sequence.

### Statistical analysis

All statistical analysis was performed in Python and R (v. 3.3.1). Cohorts (v. 0.4.0, https://github.com/hammerlab/cohorts) was used to orchestrate the analysis. The Mann Whitney and Fisher’s Exact test were performed using the Python scientific computing library, SciPy (v. 0.18.1). Kaplan-Meier curves were computed with Lifelines (v. 0.9.1.0). Survival and logistic regression models were estimated using PyStan (v. 2.12.0.0) and the Stan statistical computing software (v. 2.12.0). Survival analyses utilized a proportional hazards piecewise exponential model with a random walk baseline hazard. The analysis for presence of a time-varying covariate effect was performed in R using survival (v. 2.39.5) to look for the association of scaled Schoenfeld residuals with log(time), whereas the estimation of the time-varying covariate effect was performed using Stan. This analysis estimated the covariate effect at each timepoint with a random-walk prior. In some cases, alternative specifications of models written in Stan were interrogated as sensitivity analyses; see the project’s GitHub repository (https://github.com/hammerlab/multi-omic-urothelial-anti-pdl1) and the S1 Supplementary Analyses section for details.

All analysis code is available at https://github.com/hammerlab/multi-omic-urothelial-anti-pdl1 for open access by readers.

### Data availability

Mutation calls, TCR sequencing and RNA sequencing data are available at http://doi.org/10.5281/zenodo.546110. Additional data are available at https://github.com/hammerlab/multi-omic-urothelial-anti-pdl1.

## Results

### Patient Characteristics

29 patients with metastatic urothelial cancer from a single institution treated with atezolizumab as part of a single-arm phase II study (IMvigor 210, NCT 02108652) were included in the analyses. The patients displayed characteristics typical of the metastatic urothelial cancer population studied in IMvigor 210: 25 of 29 were males with urothelial cancers of bladder origin, and 18 of 29 had a reported prior smoking history (Table 1). Patients had an ECOG performance status of 0 or 1, and had 0 to 3 prior regimens of chemotherapy. Of this group, 25 patients had sufficient tumor tissue for WES, 26 for RNA-seq and 24 for TCR-seq. 29 had a pre-treatment peripheral blood sample on which TCR-seq could be performed; 24 had one pre-treatment and at least one post-treatment peripheral blood collection.

**Table 1:** Baseline characteristics

| Table 1: Baseline characteristics |  |  |
| --- | --- | --- |
| Variable | Durable Clinical Benefit<br>(n=9) | No Durable Benefit<br>(n=20) |
| Median Age (range) | 66 (57-74) | 72 (46-81) |
| Male Sex | 9 (100%) | 16 (80%) |
| Site of primary tumor |  |  |
| Bladder | 7 (78%) | 19 (95%) |
| Upper tract | 2 (12%) | 1 (5%) |
| ECOG performance status |  |  |
| 0 | 0 (0%) | 1 (5%) |
| 1 | 9 (100%) | 19 (95%) |
| History of tobacco use |  |  |
| No | 4 (44%) | 7 (35%) |
| Yes | 5 (56%) | 13 (65%) |
| Hemoglobin concentration < 10gm/dL | 1 (11%) | 3 (15%) |
| Albumin ≤ lower limit of normal | 2 (22%) | 5 (25%) |
| Metastatic sites at baseline |  |  |
| Visceral<sup>^</sup> | 3 (33%) | 16 (80%) |
| Liver | 2 (22%) | 9 (45%) |
| Lymph node only | 6 (67%) | 4 (20%) |
| Number of previous systemic regimens in the metastatic setting |  |  |
| 0 | 5 (56%) | 3 (20%) |
| 1 | 2 (22%) | 15 (70%) |
| 2 | 1 (11%) | 1 (5%) |
| ≥ 3 | 1 (11%) | 1 (5%) |
| Previous neoadjuvant or adjuvant chemotherapy, with first progression within ≤12 months | 3 (33%) | 2 (10%) |
| Time since previous chemotherapy ≤ 3 months | 2 (29%) | 5 (26%) |
| Prognostic risk group for previously treated patients* |  |  |
| Low | 3 (42%) | 7 (37%) |
| Intermediate | 2 (29%) | 5 (26%) |
| High | 2 (29%) | 7 (37%) |
| Prognostic risk group for previously untreated patients† |  |  |
| Low | 0 (0%) | 0 (0%) |
| Intermediate | 2 (100%) | 0 (0%) |
| High | 0 (0%) | 1 (100%) |
| Intravesical BCG administered (%) | 2 (22%) | 10 (50%) |
| ^Visceral metastasis defined as liver, lung, bone, or any non-lymph node or soft tissue metastasis.*Based on [13] †Based on [14] |  |  |

### Intratumoral and peripheral TCR features associate with durable clinical benefit

The importance of T cells to the anti-tumor response has long been known [15]; the relevance of intratumoral and peripheral T cell receptor (TCR) clonality to the anti-tumor response is an area of active study. A single previous study of melanoma patients treated with anti-PD-1 therapy demonstrated that patients whose tumors featured both high levels of tumor infiltrating T lymphocytes (TIL) along with high TIL clonality were more likely to experience radiographic response to therapy [8]. A separate study examined the peripheral TCR repertoire in anti-CTLA-4-treated patients with prostate cancer or melanoma, and found that clonotype stability was associated with response [16]. To our knowledge, no prior study has reported both intratumoral and peripheral TCR clonality in a single population treated with checkpoint blockade therapy.

We performed TCR sequencing of tumors and peripheral blood mononuclear cells (PBMC) at serial time points in our cohort. Due to limitations in sample availability (Methods), this analysis included tumors from 24 patients and peripheral blood from 29 patients, including pre-treatment samples in all patients, and between 1 and 8 total time points. A median of 141,255 (range 43,052-335,089) T cells were analyzed per peripheral blood sample, including 82,636 (range 24,095-207,860) unique TCRs, with a median clonality of 0.080 (range 0.014-0.37) and a median T cell proportion of 0.31 (range 0.082-0.64). In the tumors, the corresponding values included 1,402 (range 63-133,167) T cells, 1,086 (range 67-56,273) unique TCRs, clonality of 0.096 (range 0.033-0.34) and T cell proportion of 0.097 (range 0.0098-0.33).

In our patient group, we first asked whether there was an association between outcome and either TIL clonality or TIL proportion, or with clonality in the peripheral blood. Consistent with the data from Tumeh and colleagues [8], tumors from patients who experienced a DCB exhibited a higher TIL proportion than those patients who experienced progressive disease, with a median of 0.21 (range 0.049-0.33) in tumors from patients who had PFS greater than 6 months, versus 0.069 (range 0.0098-0.24) in tumors from patients who did not (n=24, Mann-Whitney p=0.047, Fig 1A). The consistency of result is notable given that Tumeh and colleagues used a different methodology (IHC) to assess TIL proportion than was used in our study. However, TIL proportion was not associated with continuous PFS (n=24, log-rank p=0.32) or OS (n=24, log-rank p=0.26) when split by median proportion. TIL clonality alone was not significantly associated with DCB (n=24, Mann-Whitney p=0.10, Fig 1A), continuous PFS (split by median clonality, n=24, log-rank p=0.51), or continuous OS (split by median clonality, n=24, log-rank p=0.47). Tumors with less than the median TIL proportion or TIL clonality, considered jointly as one feature, were less likely to display DCB (Fig 1B, 25% of patients with DCB versus 81% of patients without DCB, n=24, Fisher's Exact p=0.021). Considering TIL proportion and TIL clonality separately, when using median as a threshold, did not result in a significant difference in terms of DCB in either case (S1A Fig). It remains unclear whether TIL clonality adds to TIL proportion in its association with DCB in this study (TIL proportion and TIL clonality versus TIL proportion alone, n=24, log-likelihood p=0.100).

**Fig 1.**
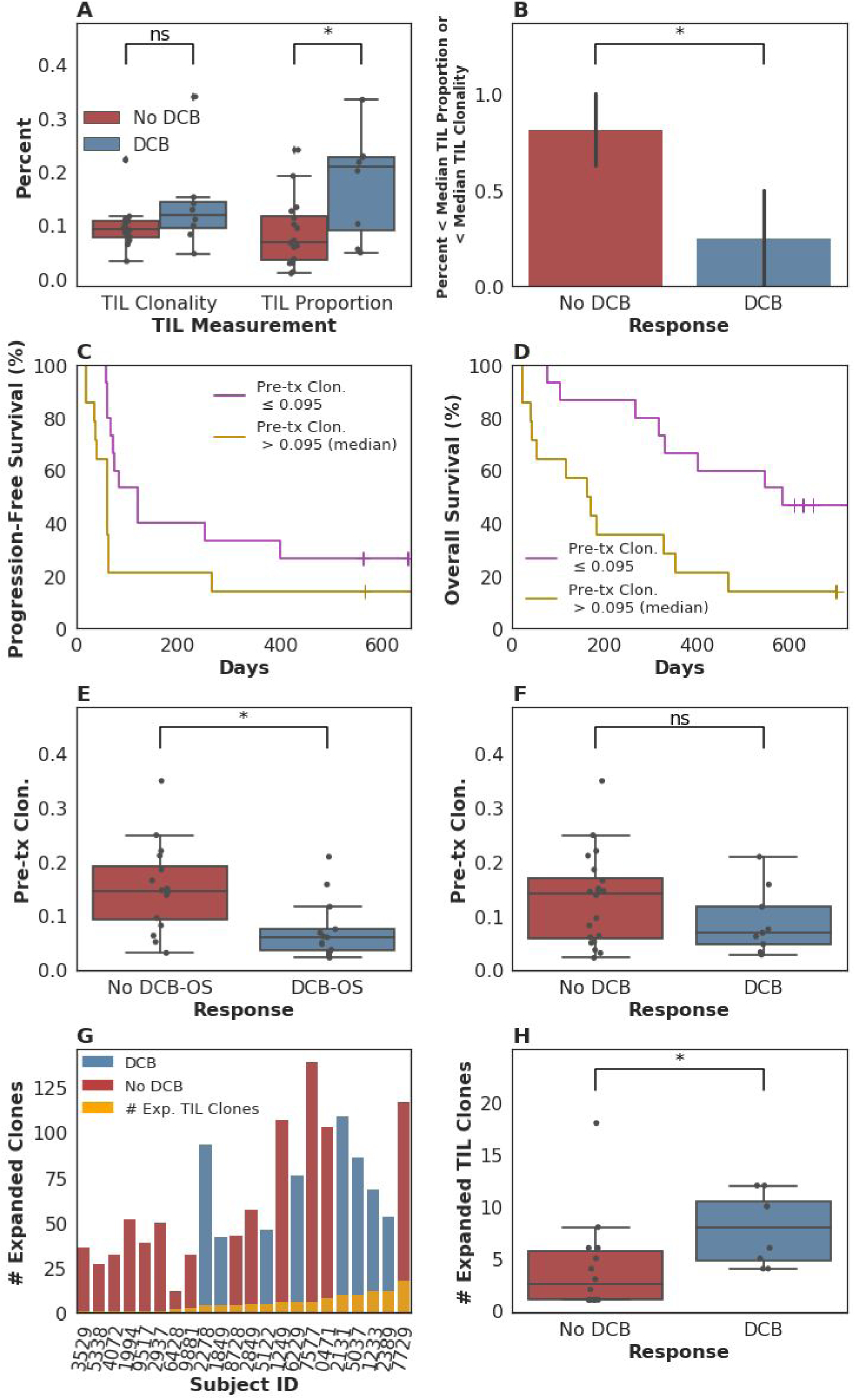
T-cell receptor clonality and treatment response. (A) TIL proportion alone was associated with DCB, with a median of 0.21 (range 0.049-0.33) in tumors from patients who had DCB, versus 0.069 (range 0.0098-0.24) in tumors from patients who did not (n=24, Mann-Whitney p=0.047). TIL clonality alone was not significantly associated with DCB, with a median of 0.12 (range 0.047-0.34) in tumors from patients with DCB and a median of 0.092 (range 0.033-0.22) in tumors from patients without DCB (n=24, Mann-Whitney p=0.10). (B) Tumors with less than the median TIL proportion or TIL clonality, considered jointly as one feature, were less likely to display DCB (Fig 1B, 25% of patients with DCB versus 81% of patients without DCB, n=24, Fisher's Exact p=0.021). (C) Patients with pre-treatment peripheral TCR clonality less than the median exhibited improved progression free survival (n=29, log-rank p=0.048). (D) Patients with pre-treatment peripheral TCR clonality less than the median exhibited improved overall survival (n=29, log-rank p=0.011). (E) There was a significant association between TCR clonality in the peripheral blood prior to initiating treatment and overall survival greater than 12 months (DCB-OS) (DCB-OS: TCR clonality 0.060 (range 0.022-0.21); OS less than 12 months: 0.15 (range 0.031-0.35) (n=29, Mann-Whitney p=0.0061)). (F) There was no significant association between pre-treatment peripheral TCR clonality and DCB (DCB: TCR clonality 0.068 (range 0.027-0.21); no DCB: 0.14 (range 0.022-0.35) (n=29, Mann-Whitney p=0.25)). (G) Expansion of TCR clones found in the tumor infiltrating T lymphocytes (TIL, orange bars) occurred in the peripheral blood 3 weeks after initiating treatment in all patients. (H) The number of TCR clones found in TIL that expanded in the peripheral blood 3 weeks after initiating treatment was 8.00 (range 4.00-12.00) in patients with DCB and 2.50 (range 1.00-18.00) in non-DCB patients (n=22, Mann-Whitney p=0.022).

We next examined pre-treatment peripheral blood clonality and its relationship to DCB. Because a diverse TCR repertoire in circulation may increase the likelihood that a tumor specific T cell population is present, we hypothesized that T cell receptor clonality would be inversely associated with response. We found that low pre-treatment peripheral TCR clonality was associated with improved PFS (split by median clonality, n=29, log-rank p=0.048), OS (split by median clonality, n=29, log-rank p=0.011), and DCB-OS (overall survival greater than one year, n=29, Mann-Whitney p=0.0061) (Figs 1C, 1D, 1E), although not with DCB (Fig 1F, n=29, Mann-Whitney p=0.25).

Finally, we explored the relationship between intratumoral and peripheral TCR clonality. Individual T cell clones present in tumors can be tracked in the peripheral blood during treatment (examples in S1B Fig). Expansion of tumor-associated TCRs occurred in the peripheral blood in all patients (Fig 1G). However, a more pronounced expansion of intratumoral TCR clones was observed in DCB patients at three weeks after initiation of treatment (second dose of therapy) (n=22, Mann-Whitney p=0.022, Fig 1H) that was not significant at 6 weeks after therapy initiation (n=20, Mann-Whitney p=0.17, S1C Fig). Interestingly, all patients with low pre-treatment peripheral TCR clonality and high TIL clonality survived greater than one year (DCB-OS, S1D Fig).

### Association of tumor genetic features with progression free or overall survival

To further examine intratumoral factors associated with therapeutic efficacy, we performed WES on 25 formalin fixed, paraffin-embedded (FFPE) archived tumor samples. Mean target coverage was 129 (range 44-194) in tumors, and 73 (range 59-91) in normal tissue. Single nucleotide variants were identified and annotated as silent, missense or nonsense mutations (Fig 2A). There was no significant association between median missense mutation load and DCB (median mutations per megabase 3.24 (range 0.038-11.46) in patients with DCB compared to 0.45 (range 0.019-9.90) in those without DCB, n=25, Mann-Whitney p=0.22, Fig 2B). There was also no significant association between missense mutation load and overall survival greater than 12 months (n=25, Mann-Whitney p=0.37, S2A Fig). In a survival analysis for time to disease progression or mortality, the estimated hazard ratio associated with increase in missense SNV count per megabase was 0.92 (95% CI 0.78 - 1.09). These results are not surprising given that the present sample size (n=25) is underpowered to detect an effect of magnitude similar to that observed by Rosenberg and colleagues [2] (power=0.2 assuming median of 12.4 vs 6.4 mutations per megabase among patients with DCB vs non-DCB response).

**Fig 2.**
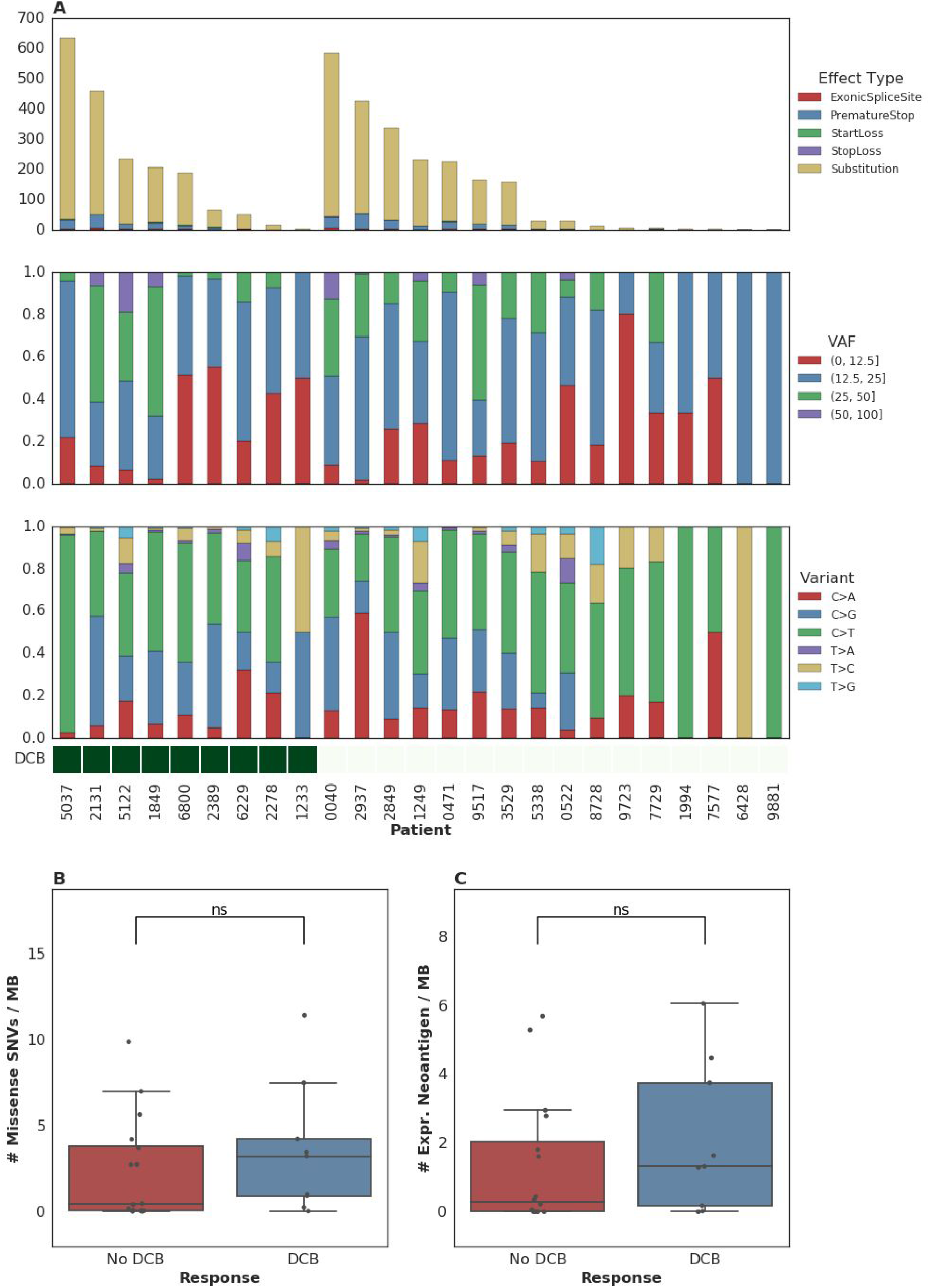
Single nucleotide variants and treatment response. (A) Single nucleotide variants, premature stop codons, transversions, mutations in start or stop codons and splice site variants as well as transitions and transversions were called for all samples. (B) Median mutations per megabase of 3.24 (range 0.038-11.46) in tumors from patients who progressed at or after 6 months, as compared to 0.45 (range 0.019-9.90) in those who progressed in less than 6 months (n=25, Mann-Whitney p=0.22). (C) Median expressed neoantigens in tumors from patients who progressed at or after 6 months was 1.32 (range 0.00-6.06), versus 0.29 (range 0.00-5.70) in those who progressed before 6 months (n=25, Mann-Whitney p=0.29).

When filtering to expressed mutations, we found a median of 0.79 (range 0.00-3.36) expressed mutations per megabase for patients with DCB and a median of 0.16 (range 0.00-3.34) expressed mutations per megabase for patients without DCB (n=25, Mann-Whitney p=0.26, S2B Fig). Consistent with the known importance of specific variant calling pipelines to output [17,18], we found that different filtering techniques impacted the association with DCB (S1 Table). Missense mutation load, when counting only mutations that were removed after post-processing (via Base Quality Score Recalibration (BQSR) and depth/variant allele frequency (VAF) filtering), was predictive of response (n=25, Mann-Whitney p=0.0078).

One hypothesis for explaining the association between mutation load and outcome to treatment with checkpoint blockade is the generation of neoantigens, altered peptides presented by the major histocompatibility complex that are capable of eliciting an anti-tumor T cell response and are more common with increased mutation load. After performing *in silico* HLA typing (Methods), we examined predicted neoantigens that are 8 to 11 amino acids in length resulting from the missense mutations of patients treated with atezolizumab. There was no significant association between predicted neoantigens per megabase and either DCB or 12 month overall survival (DCB-OS). Patients with DCB had a median 4.58 (range 0.037-39.48) predicted neoantigens per megabase while patients without DCB had 1.35 (range 0.00-20.22) (n=25, Mann-Whitney p=0.55, S2C Fig and S2D Fig). Filtering of predicted neoantigens to focus only on those expressed in RNA (Methods) also demonstrated no significant association between expressed predicted neoantigens and clinical benefit with atezolizumab (n=25, Mann-Whitney p=0.29, Fig 2C and S2B Fig). Here, too, we acknowledge the limitations in statistical power to detect associations due to the sample size of our study.

### The association between mutation load and response likelihood strengthens over time

Given that the mutation load and outcomes were weakly associated in the complete IMvigor210 dataset and not statistically significantly associated in this cohort, we embarked upon an exploration of additional factors, including tumor microenvironmental and systemic measures, which may modify the importance of this variable or independently affect outcomes.

To this end, we examined the time-varying association between mutation load and PFS to see if mutation load had a differential association with early hazards in contrast to late hazards. We found evidence of time-varying effects of somatic mutation load on progression-free survival in this cohort (n=25, p=0.044, see Methods). We estimated these effects to consist of a stronger association of somatic mutation load with mortality or disease progression more than 3 months after treatment (n=11, HR=0.69, 95% CI (0.38, 0.99)) as compared to that during the first 3 months (n=25, HR=0.91, 95% CI (0.75, 1.07); Fig 3A). This effect estimate yielded a p-value for interaction of 0.1, which does not contradict the test for presence of the effect (n=25, p=0.044) because that test is better powered. When a similar analysis was performed for time-varying association with OS, the evidence in support of the existence of time-varying effects was similar (n=25, p=0.082; notable, despite p>0.05, given the PFS results above). In terms of the estimate of these effects, patients who survived longer than 3 months exhibited a stronger association between the number of somatic mutations per megabase and a lower risk of subsequent mortality (n=11, HR=0.80, 95% CI (0.60, 1.00)) as compared to that during the first three months (n=25, HR=1.02, 95% CI (0.79, 1.22), Fig 3B). This suggests that the time-varying effect is not likely an artifact of differential association with survival versus progression. Looking at the Kaplan-Meier estimates of progression-free survival among patients with mutation load per megabase above and below the median value of 1.03, it is apparent that there is very little separation of these two populations until approximately 3 months after treatment, and that there is a high frequency of both progression and mortality events at this time (Fig 3C). While we report these results using a pre-specified threshold of 3 months, in a nonparametric analysis we found that the reduction in risk associated with somatic mutation load increased steadily over time without the emergence of a clear inflection point (S3C Fig; see S1 Supplementary Analyses and S3D Fig for further model interrogation). That said, we note that the 95% confidence intervals around the hazard ratios prior to 3 months include 1 (all p-values > 0.05; aggregate p(HR>1)=0.21) while those after 3 months are significantly less than 1 (all p-values < 0.05; aggregate p(HR>1)=0.020; p=0.07 for interaction comparing aggregate HRs). This suggests that our threshold of 3 months, which was selected based on clinical experience, may be a convenient summary of these results.

**Fig 3.**
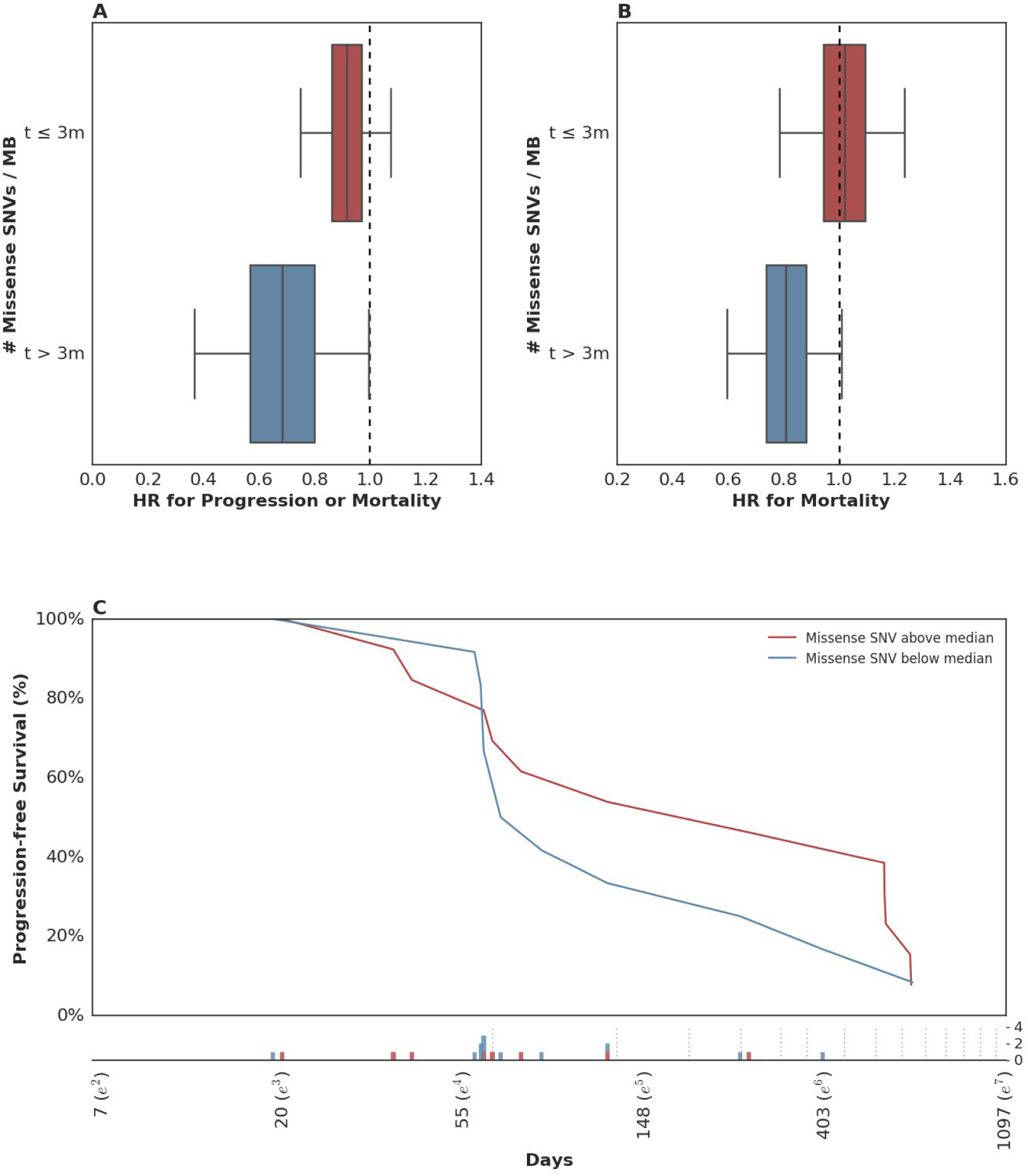
Time dependent relationship between mutation load and treatment response. (A) There was a stronger association between somatic mutation load and PFS for events occurring more than 3 months following therapy (blue box: HR=0.69, 95% CI (0.38, 0.99)), as compared to those in the first 3 months (red box: HR=0.91, 95% CI (0.75, 1.07)). (B) There was a stronger association between somatic mutation load and OS for events occurring more than 3 months following therapy (blue box: HR=0.80, 95% CI (0.60, 1.00)), as compared to those in the first 3 months (red box: HR=1.02, 95% CI (0.79, 1.22)). (C) Kaplan-Meier estimates for PFS among patients with missense SNV per megabase above the median observed value of 1.03 (red) and those with counts below the median value (blue). Time (Days) is plotted on a log-scale. For context, the frequency of observed events (progression and/or mortality) is plotted below the x-axis among patients with missense SNV per megabase above (red) and below (blue) the median value with the approximate schedule of follow-up scans per the study protocol (see Methods) shown as vertical dotted lines.

These data suggest that in patients with rapidly progressive disease, factors other than mutation load likely determine their outcome. This observation is not surprising in that clinical factor analysis of this disease state has identified a heterogeneous population of patients, with 5 clinical factors distinguishing those likely to experience a rapid and early death from those more likely to survive longer [13]. We hypothesized that such patients might simply be too clinically and systemically unwell to mount the necessary immune response, despite some of them harboring tumor biomarkers thought to confer a likelihood of DCB, including elevated mutation load. When we examined the 5-factor score in this subset relative to the rest of the dataset, we found that indeed patients who survived less than or equal to 3 months exhibited a significantly higher 5-factor score (3.00 (range 2.00-4.00), in contrast to 1.50 (range 0.00-4.00) in patients who survived longer than 3 months (n=26, Mann-Whitney p=0.018, S3A Fig). Patients surviving less than 3 months were much more likely to have liver metastases: 100% in patients surviving less than or equal to 3 months and 22% in patients surviving longer than 3 months (n=29, Fisher's Exact p=0.00097, S3B Fig). There were no significant differences in these patients with respect to BCG exposure (n=29, Fisher's Exact p=0.20), missense SNV load (n=25, Mann-Whitney p=0.26) and pre-treatment peripheral TCR clonality (n=29, Mann-Whitney p=0.12). Keeping in mind the limited sample size of this cohort, these data suggest that there is a subset of nearly end-stage patients with cancer in whom clinical variables may negate immunological response despite the presence of one or more favorable tumor-associated biomarkers. The inclusion of these clinical variables is warranted in future studies.

### Examination of the tumor microenvironment shows evidence for adaptive immunity and suppression in responding tumors

Several studies have suggested that an “inflamed” tumor microenvironment, tumor or immune cell PD-L1 expression increase the likelihood of response to checkpoint blockade. As seen in the published IMVigor 210 cohort, PD-L1 IC expression was significantly associated with DCB in this subset (n=29, Spearman rho=0.48 p=0.0083, S4A Fig). We quantified immune infiltration from RNA-seq using ESTIMATE [19]. The immune score, while associated with the TIL proportion estimated through TCR-seq (S4B Fig), was estimated to be 764.37 (range -1195.08-1509.65) in patients with DCB and 263.49 (range -1100.78-1734.28) in patients without DCB but was not significantly different (n=26, Mann-Whitney p=0.33, S4C Fig). When we performed gene set enrichment analysis (GSEA) using the Hallmark Geneset [20], we did not observe any differentially expressed gene sets between tumors from patients with DCB versus no DCB. Furthermore, RNA expression of *PD-L1* did not correlate with reported immune cell PD-L1 staining level (n=26, Spearman rho=0.045 p=0.83, S4D Fig). We did not observe a difference in tumor MHC class I expression according to DCB (S4E Fig, *HLA-A*: n=26, Mann-Whitney p=0.26, *HLA-B*: n=26, Mann-Whitney p=0.36, *HLA-C*: n=26, Mann-Whitney p=0.24).

Given that such agnostic approaches did not reveal a clear association between tumor microenvironment factors and response, we pursued a hypothesis-driven approach examining the genes that show upregulation at the cell surface during T cell exhaustion. When categorized by DCB, there was no significant difference in expression of such genes, including *CTLA-4, TIGIT, HAVCR2 (TIM-3)* or *LAG-3* [21]. When grouped by PD-L1 staining, we found low expression of all markers in the PD-L1 low group (IC0), as expected. However, in the PD-L1 high group (IC2), *HAVCR* exhibited significantly higher expression in tumors from patients who experienced DCB than in those who did not (S4F Fig). Interestingly, of the three IC2 tumors, two had missense SNV loads significantly below the median (17 and 57); the third had 412 SNVs. Additionally, although Rosenberg and colleagues [2] found that among the four TCGA subtypes of RNA expression, luminal cluster II showed a significantly higher response rate, no significant association was found here between the four clusters and DCB (n=20, Fisher's Exact p=0.36) (S4G Fig), nor between the luminal/basal sub-categorization and DCB (n=20, Fisher's Exact p=1.00), possibly due to sample size.

### Relative importance of somatic, immune, and clinical factors in resistance and response to PD-L1 blockade

Unanswered questions that arise from the many studies of biomarker correlates of checkpoint blockade response are whether measures such as mutation load, PD-L1 staining, and others reflect the same “tumor state,” or if each confers an independent effect on outcome?

When examined in conjunction with mutation load, the greater the expression of PD-L1, the more negative the association of mutation load with hazard (i.e. higher mutation load was associated with longer survival). Among patients with tumors showing little-to-no expression of PD-L1 (IC0 rated), each unit increase in missense SNV count per megabase was associated with a negligible change in hazard (n=4, HR=1.43, 95% CI (0.75, 2.98)). Among patients with tumors expressing PD-L1 at moderate or high levels (IC1 or IC2 staining), missense SNV count per megabase was associated with lower risk for disease progression or mortality (among IC1: n=11, HR=0.75, 95% CI (0.47, 1.14); among IC2: n=10, HR=0.73, 95% CI (0.48, 1.06)). Although our limited sample size precludes making an assertion that mutation load is associated with survival in any particular subgroup (e.g. when looking among IC1 and IC2 tumors alone); our data do support the presence of an interaction among these variables (p=0.046 for interaction; S5A Fig). Given the plausibility of the finding that somatic mutation load may correlate better with survival among patients with an inflamed tumor microenvironment, the addition of somatic mutation load to PD-L1 IC staining warrants further study.

We found a similar albeit weaker interaction effect when looking at association of somatic mutation load (missense SNV count per megabase) and progression-free survival according to the presence/absence of liver metastasis prior to treatment administration (p=0.14 for interaction). Among patients without liver metastasis, somatic mutation load was associated with a lower risk for disease progression or mortality (n=16, HR=0.73, 95% CI (0.50, 1.02), S5B Fig) than patients with liver metastasis (n=9, HR=0.96, 95% CI (0.66, 1.37), S5B Fig).

To our surprise, although both PD-L1 staining and mutation load were each associated with response in the original study [2] these variables did not correlate with each other (Fig 4A). Furthermore, pre-treatment peripheral TCR clonality did not correlate with mutation load (Fig 4B). The lack of association between these variables suggests that each might have an independent or semi-independent role in determining the likelihood of response to therapy. TCR clonality and infiltration did, however, correlate with PD-L1 IC score: those tumors with higher clonality or higher infiltration also featured higher PD-L1 staining (p=0.02 and p=0.01, respectively, Figs 4C, 4D).

**Fig 4.**
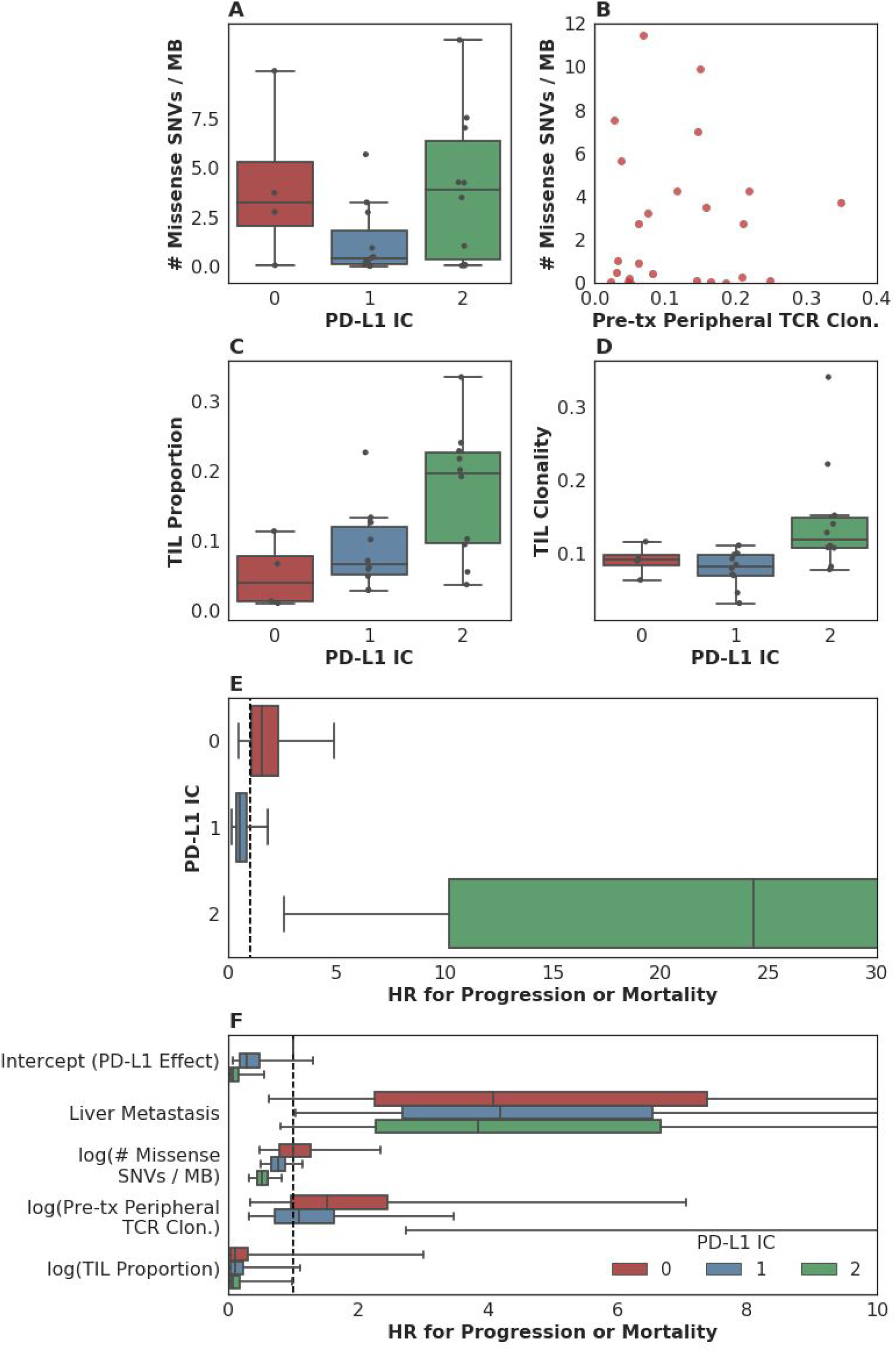
Associations between measured somatic, immune and clinical variables. (A) Although both PD-L1 staining and mutation load each were weakly associated with response, these variables were not correlated with each other (n=25, Spearman rho=0.14 p=0.51). (B) Pre-treatment peripheral TCR clonality did not correlate with mutation load (n=25, Pearson r=0.0017 p=0.99). (C) TIL proportion as estimated by TCR sequencing was associated with PD-L1 IC staining (n=24, Spearman rho=0.51 p=0.010). (D) TIL clonality was associated with PD-L1 IC staining (n=24, Spearman rho=0.48 p=0.017). (E) Hazard associated with log(pre-treatment peripheral TCR clonality) by level of immune cell PD-L1 expression (IC0, IC1 or IC2). (F) Multivariate survival analysis of various clinical, peripheral and intratumoral biomarkers for association with time to disease progression or mortality (PFS), utilizing a varying-coefficient model which allows the hazard associated with a one-unit increase in a biomarker’s value to vary according to level of intratumoral PD-L1 expression (IC score).

In an analysis to see whether the association between pre-treatment peripheral TCR clonality and progression-free survival varied by PD-L1 IC score, we found some evidence of an interaction (p=0.015 for interaction; Fig 4E). Among patients with low levels of PD-L1 expression, there was little association between pre-treatment peripheral TCR clonality and progression-free survival (among IC0: n=4, HR (mean)=1.86, HR (median)=1.55, 95% CI (0.50, 4.99), p(HR>1)=0.21; among IC1: n=11, HR (mean)=0.69, HR (median)=0.58, 95% CI (0.15, 1.84), p(HR<1)=0.19)). Among patients with high levels of PD-L1 expression, by comparison, we observed almost complete separation of progression-free survival according to pre-treatment peripheral TCR clonality (among IC2: n=10, HR (mean)=89.88, HR (median)=23.41, 95% CI (2.43, 506.94), p(HR>1)=0.0014; Fig 4E). Similar results were seen in analyses with respect to OS, and in a logistic regression analysis for DCB (S5C Fig, S5D Fig, S5E Fig).

To resolve the hypothesis that those patients with low peripheral TCR clonality simply were healthier, we examined the association between 5-factor score and pre-treatment peripheral TCR clonality and did not find such an association (n=26, Spearman rho=0.25 p=0.22, S5F Fig).

In a multivariate survival model for time to disease progression or mortality, which allows the effect of each biomarker to vary according to intratumoral PD-L1 IC score, we find that the correlation of each intratumoral, peripheral or clinical biomarker with disease progression or mortality is relatively independent of the others (Fig 4F, S5G Fig). Perhaps with the notable exception of the association between liver metastatic status and time to progression or survival, the correlation of each intratumoral or peripheral biomarker with outcome is strongest in the group with the highest levels of immune cell PD-L1 expression (S2 Table).

## Discussion

The treatment of previously incurable metastatic solid tumors with checkpoint blockade agents has led to dramatic success in a minority of patients, a finding that has generated substantial excitement in the field, with associated correlative studies and drug development. Here, we undertook the in-depth characterization of tumors and peripheral blood from 29 patients treated on IMvigor 210, a Phase II study in which 310 patients were treated with the anti-PD-L1 agent atezolizumab. In this cohort of patients, we illustrate the importance of host immune factors, including intratumoral and peripheral T cell receptor clonality, infiltration and expansion, to clinical outcomes. We did not find a significant association between mutation or expressed neoantigen load and progression free survival or DCB (defined as progression free survival (PFS) > 6 months). However, we did observe a time-dependent relationship between mutation load and outcome, wherein a relationship between mutation burden and outcome could only be detected in those patients surviving greater than 3 months. This analysis implies that patients who experienced rapid progression may display systemic indicators of immune deficiency despite elevated mutation load in the tumors. Calculation of the hazard ratios for each measured biomarker and clinical factor underscores the concept that a complex interaction of both host and tumor variables determines whether a patient will experience clinical benefit from anti-PD-L1 therapy. Although the overall study found significant associations between mutation load as measured by the Foundation Medicine targeted sequencing panel and radiographic response [2], there was no statistically significant association between mutation load and durable clinical benefit or survival in the patient subset studied here, despite the similarity of our study population to the parent study. This contrast may be due to a combination of factors. First, though statistically significant, the association in the overall study was not categorical: as in other studies of mutation load, this factor alone was not predictive of response. Second, we have less power to detect this association in our smaller subset compared with the larger studied cohort. Third, standardized definitions and calculations of mutation load do not exist as yet; each published study has used differing methodologies [2,10,22,23]. Indeed, in this study, depending on the method used, the association between mutation load and clinical outcomes varied from p<0.08 to p>0.4 (AUCs and p-values in S1 Table). To illustrate the fickle nature of defining mutation load, counting only the mutations excluded by BQSR, as opposed to only those remaining after BQSR, showed a significant association with DCB. Together, these findings underscore the need for improved and standardized mutation calling methods. The weak association of mutation load with DCB and the lack of such standardization render this biomarker unfit for application to individual patients at present. Furthermore, if validated in another dataset, this analysis implies that a clinical and immunological state may exist in patients with advanced cancer, such that patients with very rapidly progressing disease and expected death in less than 3 months do not respond despite the presence of positive intratumoral biomarkers.

In an attempt to deepen our understanding of the biology of response and resistance, we studied additional factors. We found that even in this small dataset, TCR clonality below the median in the peripheral blood prior to treatment, expansion of tumor-associated TCR in the periphery 3 weeks after initiating treatment, and higher TIL proportion are all associated with clinical benefit. These data suggest that TCR sequencing provided additional insights into response and resistance beyond mutation load and PD-L1 staining. With respect to biomarker development, our study implies that non-invasive metrics such as pre-treatment peripheral TCR clonality and known prognostic features such as the presence of liver metastases may be worthy of further study in urothelial cancer patients treated with PD-L1 blockade.

From a mechanistic perspective, these findings imply an important relationship between circulating and intratumoral immunity upon PD-L1 blockade. We hypothesize that low TCR clonality in the peripheral blood prior to treatment increases the likelihood that a patient harbors one or several clones capable of tumor recognition, whether of neoantigens or tumor-associated antigens. The expansion of tumor-associated TCRs in the peripheral blood underscores the continuity of the tumor and blood compartments, and suggests that the activity of PD-L1 blockade may involve circulating T cells more than was previously thought. Indeed, this raises the possibility that anti-tumor T cells may home from the periphery to the tumor before later recirculating.

Finally, though limited in power by the small sample size, we attempted to integrate the importance of the studied variables. This analysis demonstrated both hypothesized and unexpected interactions. For example, while mutation load seemed to be associated with outcome more significantly in PD-L1 IC1 and IC2 tumors, high PD-L1 IC staining in the setting of high peripheral TCR clonality was associated with a substantial hazard for poor outcome. Given the significance of PD-L1 expression in mediating response to anti-PD-L1 therapy, the presence of these interactions may argue in their favor as predictive rather than prognostic biomarkers. Further analysis is required to elucidate the role of these biomarkers in mediating response to checkpoint blockade.

This study has several limitations. The patients under study were treated at a single institution and represent a small subset of the overall study, limiting statistical power. As a single arm Phase II study, there is no control arm for comparison. Tumor samples were FFPE and were not collected immediately prior to treatment initiation. Only one sample per patient was utilized, which does not necessarily capture the heterogeneity of each tumor. Finally, a number of analyses were performed on a small number of patients without an independent validation cohort; although most analyses were pre-specified, there is a risk of Type II error and no adjustments were made for multiple testing.

In conclusion, we have demonstrated the potential value of pursuing an integrated study of somatic, immune and clinical features in order to elucidate the biological mechanisms underlying response to checkpoint blockade and ultimately improve our ability to practice precision medicine. We hope this work will motivate further multi-omics studies of response to checkpoint blockade.

## Acknowledgements

Carol Aghajanian and Jedd Wolchok for their mentorship of AS.

## Supplementary Information, Captions

### Supplementary Tables

S1 Table. Choice of depth and VAF filtering, as well as whether or not to run Base Quality Score Recalibration (BQSR), resulted in differences in predictive value for mutation load. We chose to optimize for precision, choosing the highlighted set of filters/BQSR. Precision is defined as the fraction of filtered missense mutations in IMPACT genes that were actual IMPACT panel variants. Recall is defined as the fraction of actual IMPACT panel variants that were found in the filtered missense results.

S2 Table. Summary of results from multivariate survival analysis of various clinical, peripheral and intratumoral factors to estimate their independent association with hazard for disease progression or mortality (PFS) and for mortality (OS) according to level of intratumoral PD-L1 expression (IC grade). Results are summarized as median and 95% posterior intervals.

Supplementary Figures

S1 Fig. (A) Fig. 25% of patients with DCB had less than the median TIL proportion versus 63% of patients without DCB (n=24, Fisher's Exact p=0.19); similarly, 25% of patients with DCB had less than the median TIL clonality versus 63% of patients without DCB (n=24, Fisher's Exact p=0.19). (B) TCR overlap between the pre-treatment and 3-week posttreatment peripheral blood in one patient with limited clinical benefit (PFS=37 days) and one patient with durable clinical benefit (CR at 630 days after starting treatment). The association between pre-treatment peripheral blood TCR sequences (x axis) and posttreatment peripheral blood TCR sequences (y axis) is overlaid with the presence of tumor associated T cell clones. Gray indicates TCRs present only in the peripheral blood; blue indicates TCRs present in the tumor and blood; orange indicates TCRs present in the tumor and expanded in the blood with treatment. (C) There was no significant expansion of TIL-associated TCR clones between pre-treatment (3.00 (range 1.00-9.00)) and 6 weeks post-treatment (2.00 (range 1.00-8.00)), n=20, Mann-Whitney p=0.17. (D) The combination of high pre-treatment TIL and low pre-treatment peripheral blood TCR clonality were predictive of DCB (n=24, Fisher's Exact p=0.0069) and DCB-OS (n=24, Fisher's Exact p=0.014). For DCB, a logit model combining both was more predictive than peripheral blood (n=24, log-likelihood p=0.00029) or TIL (n=24, log-likelihood p=0.00051) clonality alone. For DCB-OS, both combined were more predictive than TIL (n=24, log-likelihood p=0.0029) clonality alone.

S2 Fig. (A) No significant association between the number of missense SNV per megabase and overall survival, with 2.13 (range 0.038-11.46) in tumors from those patients who survived greater than 12 months, versus 0.48 (range 0.019-9.90) in those who did not (n=25, Mann-Whitney p=0.37). (B) No significant difference between median expressed neoantigens in tumors from patients who survived greater than or equal to 12 months: was 1.31 (range 0.00-6.06), versus 0.35 (range 0.00-5.30) in those who survived less than 12 months (n=25, Mann-Whitney p=0.36). (C) No significant difference between median predicted neoantigens per megabase: 4.58 (range 0.037-39.48) in tumors from patients with DCB, as compared to 1.35 (range 0.00-20.22) in those who progressed in less than 6 months (no DCB) (n=25, Mann-Whitney p=0.55). (D) No significant difference between median predicted neoantigens per megabase in tumors from those patients who survived greater than 12 months (here used to define DCB-OS) was 3.56 (range 0.037-39.48) as compared to 1.37 (range 0.00-20.22) in those who did not (no DCB) (n=25, Mann-Whitney p=0.81).

S3 Fig. (A) Patients who survived less than 3 months (red box) exhibited a significantly higher 5-factor score (3.00 (range 2.00-4.00), as compared to 1.50 (range 0.00-4.00) in patients who survived >3mo (blue box) (n=26, Mann-Whitney p=0.018). (B) Patients who survived less than or equal to 3 months (red box) were more likely to have liver metastases (100% in patients who survived less than or equal to 3 months and 22% in patients who survived longer than 3 months, n=29, Fisher's Exact p=0.00097). (C) The hazard ratio for each mutation per megabase, estimated at each unique failure time. Red box plots summarize 50% and 95% posterior intervals for each observed failure/censor time, with median values shown in green. Time (Days) is plotted on a log-scale. Estimates are not independent from one another since the model utilizes a random-walk parameterization to allow the variance in hazard over time to be modeled flexibly. (D) Posterior predicted intervals for PFS drawn from the survival model estimating the time-varying effect of mutation count on PFS. Intervals are shown for patients with missense SNV per megabase above the median and those with counts below the median value (blue) for illustrative purposes. This cutpoint was not used in the model; missense SNV per megabase was included as a continuous covariate. Lines are drawn at median values of the posterior predictive distribution, with 50% credible intervals shown in the shaded regions. Time (Days) is plotted on a log-scale.

S4 Fig. (A) PD-L1 IC staining as reported by the sponsor in the published study [2] and outcome in our cohort were significantly associated in this sub-study (n=29, Spearman rho=0.48 p=0.0083). (B) ImmuneScore was associated with TIL proportion (n=24, Spearman rho=0.47 p=0.022). (C) There was no association between ImmuneScore and DCB (DCB, 764.37 (range -1195.08-1509.65); no DCB 263.49 (range -1100.78-1734.28) (n=26, Mann-Whitney p=0.33). (D) PD-L1 expression as measured by RNA-seq was not associated with PD-L1 IC level (n=26, Spearman rho=0.045 p=0.83). Tumor cell PD-L1 staining was not available. (E) HLA Class I expression was not associated with DCB (HLA-A: n=26, Mann-Whitney p=0.26, HLA-B: n=26, Mann-Whitney p=0.36, HLA-C: n=26, Mann-Whitney p=0.24). (F) Expression of other inhibitory markers, in particular *HAVCR2* (also known as *TIM-3*) was higher in DCB patients in the IC2 group. (G) No association was found between TCGA RNA Subtype and response in this sub-study (n=20, Fisher's Exact p=0.36).

S5 Fig. (A) Hazard associated with log(missense SNV count per megabase) by level of immune cell (IC0, IC1 or IC2) PD-L1 expression. (B) Hazard associated with log(missense SNV count per megabase) by presence or absence of liver metastasis at enrollment. (C) Association of peripheral TCR clonality prior to treatment with time to mortality (OS) varies according to immune cell (IC0, IC1 or IC2) PD-L1 expression. (D) Association of peripheral TCR clonality prior to treatment with DCB varies according to immune cell (IC0, IC1 or IC2) PD-L1 expression. (E) Association of peripheral TCR clonality prior to treatment with DCB (OS) varies according to immune cell (IC0, IC1 or IC2) PD-L1 expression. (F) There was no significant relationship between 5-Factor score and pre-treatment TCR clonality (n=26, Spearman rho=0.25 p=0.22). (G) Multivariate survival analysis of various clinical, peripheral and intratumoral biomarkers for association with time to mortality (OS), utilizing a varying-coefficient model which allows the hazard associated with a one-unit increase in a biomarker’s value to vary according to level of intratumoral PD-L1 expression (IC score). Note that the x-axis has been truncated at a value of 10 for clarity even though this results in the exclusion of some estimated HR values (specifically that for pre-treatment peripheral TCR clonality among IC2 patients).

S6 Fig. (A) No significant association between the number of missense SNV found on MSK-IMPACT and DCB (DCB 0.13 (range 0.00-0.31) versus no DCB 0.046 (range 0.00-0.37) (n=25, Mann-Whitney p=0.42). (B) No significant association between the number of missense SNV found on MSK-IMPACT and overall survival (survival greater than 12 months 0.093 (range 0.00-0.31), versus less than 12 months 0.074 (range 0.00-0.37) (n=25, Mann-Whitney p=0.78). (C) There was no significant difference in APOBEC signature found in tumors from patients with PFS DCB (0.19 (range 0.00-0.56)) as compared to no DCB (0.00 (range 0.00-0.46)) (n=25, Mann-Whitney p=0.23). (D) There was a significant correlation between missense SNV count and APOBEC signature mutations (n=25, Pearson r=0.40 p=0.048). (E) There was no significant association between FGFR3 mutations or expression (n=26, Mann-Whitney p=0.39). (F) There was no significant association between MYC expression and outcome measured by PFS DCB (n=26, Mann-Whitney p=0.87). (G) There was no significant association between DNA damage response (DDR) mutations rated as “possible” by PolyPhen and DCB (n=25, Mann-Whitney p=0.20). (H) Univariate association of exonic SNV, missense SNV and neoepitope load with DCB, with (blue bars) and without (red bars) filtering by expression. (I) Univariate association of exonic SNV, missense SNV and neoepitope load with OS greater than 12 months, with (blue bars) and without (red bars) filtering by expression. (J) Univariate association of exonic SNV, missense SNV and neoepitope load with PFS, showing results with (blue bars) and without (red bars) filtering by expression. (K) Univariate association of exonic SNV, missense SNV and neoepitope load with OS, showing results with (blue bars) and without (red bars) filtering by expression. (L) Univariate association of expressed/total ratio for exonic SNV, missense SNV, and neoantigen loads with PFS. (M) Univariate association of expressed/total ratio for exonic SNV, missense SNV, and neoantigen loads with OS. (N) Univariate association of expressed/total ratio for exonic SNV, missense SNV, and neoantigen loads with DCB (PFS greater than 12 months). (O) Univariate association of expressed/total ratio for exonic SNV, missense SNV, and neoantigen loads with OS greater than 12 months. (P) There was no significant association between pack years of reported smoking history and DCB (n=29, Mann-Whitney p=0.87). (Q) There were significant associations between PFS (n=29, log-rank p=0.024) and OS (n=29, log-rank p=0.018) and the presence of liver metastasis. ® There was a significant association between the presence of visceral metastases and poor overall survival (n=29, log-rank p=0.020). (S) There was not a significant association between 5-factor score and OS (n=26, log-rank p=0.13). (T) No significant association between the number of expressed missense SNV per megabase and DCB (DCB 0.79 (range 0.00-3.36), versus no DCB: 0.16 (range 0.00-3.34)), n=25, Mann-Whitney p=0.26.

